# Organs-on-a-Chip Database (OOCDB): A Comprehensive, Systematic and Real-time Organs-on-a-chip Database

**DOI:** 10.1101/2022.07.05.498623

**Authors:** Jian Li, Weicheng Liang, Zaozao Chen, Xingyu Li, Pan Gu, Anna Liu, Pin Chen, Qiwei Li, Xueyin Mei, Jing Yang, Jun Liu, Lincao Jiang, Zhongze Gu

## Abstract

Organs-on-a-Chip is a microfluidic microphysiological system that uses microfluidic technology to make high-resolution and real-time imaging analysis on the structure and function of living human cells at the level of tissue and organ in vitro. Compared with the traditional two-dimensional cell culture model and animal model, organs-on-a-chip technology can simulate the pathological and toxicological interactions between different organs or tissues more closely and reflect the collaborative response of multiple organs to drugs. Although lots of organs-on-a-chip-related literature have been published, none of current databases have achieved all the following functionalities yet: searching, downloading and analyzing data and results from literature of organs-on-a-chip. To address this need, we established a database named organs-on-a-chip database (OOCDB), as a platform to integrate information related to organs-on-a-chip from various sources: literature, patents, microarray and transcriptome sequencing raw data, many open access data of organs-on-a-chip and organoids, as well as the data generated in our lab. OOCDB comprises dozens of sub databases and analysis tools and each sub database contains a number of data related to organs-on-a-chip, aiming to provide a comprehensive, systematic and convenient search engine for researchers. In addition, it provides functions such as mathematical modeling, three-dimensional model and citation map to meet the needs of researchers and to promote the development of organs-on-a-chip. The organs-on-a-chip database can be visited at http://www.organchip.cn.

## Introduction

Traditional clinical drug research usually uses two-dimensional cell culture models [1, 2] and animal models [3]. However, there is structural difference between monolayer cells *in vitro* and natural tissues. The traditional *in vitro* cell culture models are also limited in their ability to compare the interactions between different organs and tissues [1,4,5]. Therefore, the efficacy of drug screening cannot be accurately tested [6, 7]. Moreover, due to the interspecific differences between human and animals, many drugs that pass the screening of animal models, may fail in clinical trials because of their toxicity to humans or little curative effect [8]. Therefore, there is an urgent need for a new technology to complete the construction of pathological and physiological models *in vitro* and organs-on-a-chip technology came into being. Organs-on-a-chip is an *in vitro* microfluidic microphysiological system [4, 9] that simulates structural and functional characteristics of human tissues and organs. It is established on microfluidic chip by using micro processing technology, cell culture, biomaterials, and other methods [10, 11]. It can not only perform high-resolution and real-time imaging analysis on the structure and function of living human cells at the level of tissue and organ *in vitro* [11], but also can eliminate the uncertainty caused by complex metabolism in traditional pharmacokinetic studies [12, 13].

In recent years, a variety of organs-on-a-chip technologies have been developed [14]. These organ-on-chips are composed of different organ specific cell types [4], which can be used to simulate corresponding organs. Some examples include lung chips [15, 16], liver chips [17], cornea chips [10], heart chips [18], *etc*. Due to the development of organs-on-a-chip technology, there is an explosive growth in disease modeling data of different organs. In addition, the complexity of human organs-on-a-chip structure, measurement connotation and the rapid development of big data also hinder the application of organs-on-a-chip data.

Among established databases, MPS-DB database of University of Pittsburgh is the only database specific to organs-on-a-chip [19, 20]. It is successful in collecting and using organs-on-a-chip data and contributes greatly to development of related database [21]. However, there are some shortcomings in this database. Firstly, MPS-DB contains a large amount of organ-on-a-chip model data, and the collection of organ-on-a-chip data is comprehensive. However, for organ-on-a-chip related data, such as literature, patents and other public data content is relatively lacking. Thus, the richness of data is not enough. Secondly, the data of MPS-DB is not fully downloadable, only the brief data of the models are available for download, but the pictures and detailed descriptions of the models are not available for download, so the availability of the data is insufficient. In addition, there are some public databases that can be used for organs-on-a-chip research, such as PubChem [22], NCBI [23], Drugbank, CHEMBL, BRD, *etc*. But due to their lacks of pertinence and standardized methods and management system for multi-dimensional and heterogeneous organs-on-a-chip big data, they are not helpful enough to research of organs-on-a-chip. Driven by the need to better prepare for future research and provide a comprehensive and searchable database for scholars in relevant fields, we have constructed the OOCDB (organs-on-a-chip database).

The OOCDB is a standardized system with a large number of charts, literature and patent information. In addition, the content of our database is updated periodically. Relevant literature, patents and other information are collected every week to inform researchers of the latest progress and research hotspots. Moreover, compared with other databases, OOCDB has the following advantages. (1) Our database is separated into sections such as literature, patents, drugs, poisons, chemicals and other information and each section contains a number of open access data. The database content is comprehensive and the data is detailed. In addition, there is a large amount of lab-generated data in the OOCDB, which complements the public data and together constitute the data source of the OOCDB. (2) Our database handles organs-on-a-chip related data, analyzes the author, publishing time and publishing unit of the data, and provides researchers with analysis of the latest research results. (3) Our database can also provide free online tools for researchers, such as mathematical modeling allows researchers to upload their own data, or data of interest to them. These data will be analyzed by tissue enrichment analysis to determine the organ of gene enrichment and thus correspond to their own experimental results and provide a reference for research. Therefore, it is of great significance to further promote the data exchange and sharing of organs-on-a-chip to accelerate the research on organs-on-a-chip for conquering complex diseases.

## The methods of the database and contents

### Data collection and integration

As shown in **Figure 1**, it is the main framework, interaction and logic of our organs-on-a-chip database. In order to be able to build a high-quality organs-on-a-chip database, OOCDB follows a standardized process for data collection and organization. As shown in Figure 1A, we use “Organ-on-a-Chip”, “Organs-on-a-Chip ”, “Organ on chips”, “Organ chips”, “Organoid chip”. “ Heart chip”, “Liver chip”, “ Vascular chip”, “Skin chip”, “Lung chip”, “Intestinal chip”, “Neural chip”, “Tumor chip”, “Reproductive System and Embryo Chip” keywords retrieved from PubMed, NCBI, BRD, Google Patent Related publications. The data were filtered by manually reading the abstracts and full text, and the data from publications not related to organ chips were excluded. A total of 107,873 relevant literature data, 16,841 histology data, and 12,002 patent data were collected. We extracted drug, toxin, compound, and device information from the literature, patent and histology data. We obtained the relevant open access data from Drugbank, ChEMBL, CTD, EMA according to keywords, and provided links to the original websites for non-open access data. A total of 14020 drug data, 17070 poison data, 4986258 compound data and 160 device data were collected.

**Figure 1.**
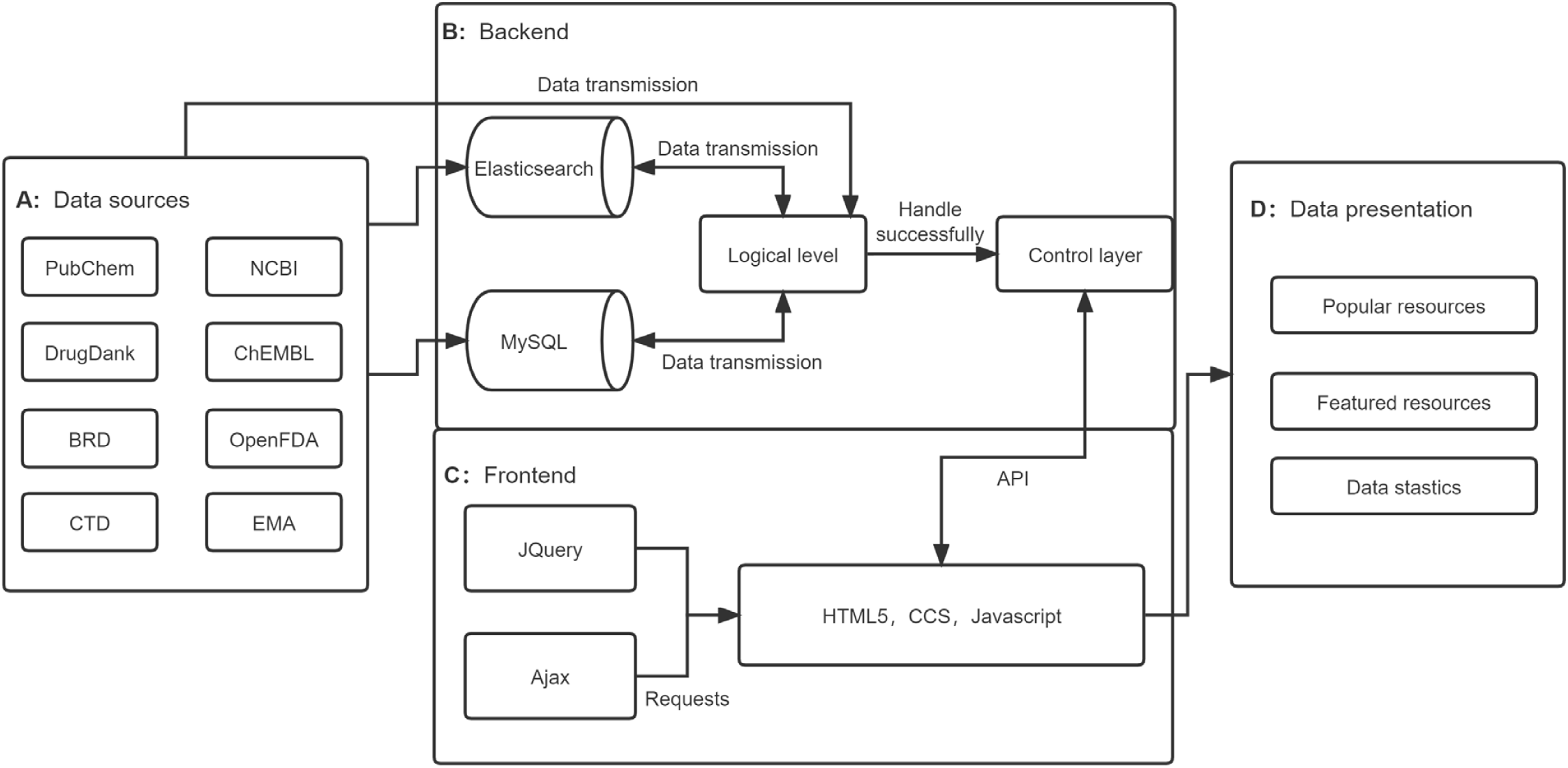
The main construction principle of organs-on-a-chip database. **A**. The data sources of the OOCDB. Mainly includes Pubchem, NCBI, DrugBank, ChEMBL, BRD, OpenFDA, CTD, EMA. **B**. The Backend of the OOCDB. The data is processed by Elasticsearch and MySQL according to the logical level, and then transferred to the control layer after processing. **C**. The Fronted of the OOCDB. Backend data transfer through the API combined with JQuery and ajax through HTML5, CCS, Javascript to write front-end web pages. **D**. The data presentation of the OOCDB. After the frontend processing, the data is mainly divided into Popular resources, Featured tools and Data Statics.

As shown in Figure 1B, we imported the collected data into the control layer for analysis and integration. For the literature data, the data were classified and processed according to different organs and different cancers studied, and put into the organ database and cancer research database, respectively. At the same time, the authors and the citation and cited relationships of the literature are processed to form the citation data. We use the citation data to form a Citation Map, which integrates more than 100,000 papers and makes the papers interrelated. In addition, we use the Elasticsearch tool to connect the literature and drug libraries, and filter out the drugs that appear in the literature and link them to the drugs in the drug library, so that the drug data can be accessed directly through the literature and the literature data can be accessed through the drugs. For patent data, we categorize them by country and count the application date and expiry date of patents, providing the option to filter patents by date. For histology data, we sort the statistics by the organ they were studied in and provide a distribution map of the organs studied. We also use the Elasticsearch tool to process the histology data and integrate it with the literature data. Finally, based on the MySQL database system, the databases in OOCDB become an interrelated system rather than separate databases.

### Database construction

We use Elasticsearch tool to build the database index. At the same time, for the synonym problem that occurs during the retrieval process, we use the multi-field index method in Elasticsearch. We set different weights in different fields so that the search weight of synonyms is less than that of the original word, ensuring that the original word is ranked first in the search results, and at the same time ensuring that synonyms can take effect in time. For the possible duplicate data, we set the MySQL database primary key to set the article title, patent title, drug name, poison name and other data as the PRIMARY KEY, which avoids the duplicate data because the database primary key is unique. As shown in Figure 1C, the data processed by the control layer is transferred to the front-end through API. The database uses jQuery framework, HTML5, CSS and JavaScript technologies for web page construction, and requests are handled through Ajax. Finally, as shown in Figure 1D, the OOCDB database, including popular resources, featured tools, and publication statistics, is formed.

### Database interface

The architecture of OOCDB is shown in **Figure 2**. In the OOCDB, there are 6 navigation bars, 8 sub-databases and 3 featured tools for data analysis processing and related statistical analysis, which can provide data related to organs-on-a-chip literature and patent data, brief information of drugs, toxins, compounds and their applications in organs-on-a-chip. As shown in Figure 2, the six navigation bar entries of the website are home page, database page, submission page, about page, help page and lab data page. The main part of OOCDB is the database page, which mainly includes popular resources, featured tools and publication statistics. The popular resources mainly include eight sub databases: GSA (Genome Sequence Archive), drug bank, device, organs, patents, cancer research, chemicals and toxicants. Each sub database has its own special data content. Featured tools achieved innovative advancement such as three-dimensional model, mathematical modeling and citation map, each of which performs specialized function. The top authors, top affiliation and top journals section in the publication statistics which is a reflection of trending research topics and latest discoveries.

**Figure 2.**
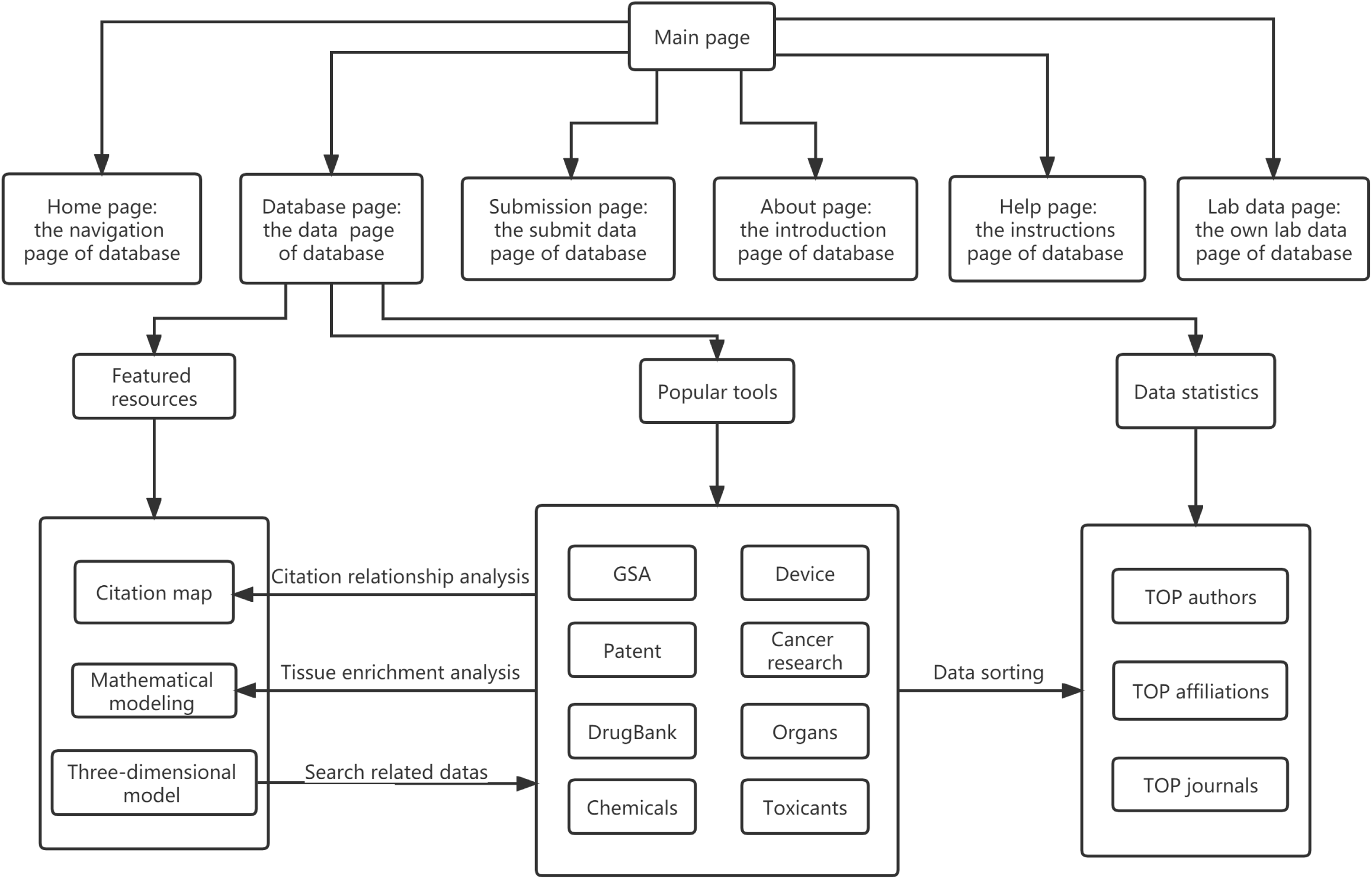
The main framework of organs-on-a-chip database: The 6 navigation bars of OOCDB are Home page, Database page, Submission page, About page, Help page and Lab data page. The main part of OOCDB is the Database page. The featured tools, popular resources and publication statistics are interrelated. The popular resources mainly store the basic data, and the featured tools and publication statistics are mainly based on the popular resources for analysis and processing.

Following information can be found through simple search in OOCDB. (1) Relevant data on organs-on-a-chip, including literature, patents and histology data. (2) The information about drugs, devices, organs, patents, chemicals, toxicants, data form the lab itself and other Organs-on-a-chip related data. (3) Names of famous experts, institutions and journals in the field of organs-on-a-chip which can be used for hot spot analysis in the field of organs-on-a-chip. In addition to easy access to the required information, OOCDB provides a variety of online tools. For example, users can use mathematical models for tissue enrichment analysis, select the database’s own standard database or a database provided by the user for comparison, and thus find the organs enriched by the genes of interest. In addition, in order to collect more organs-on-a-chip related data, users can submit their organs-on-a-chip related research results in our submission interface to enrich the database content.

## Results

OOCDB page is mainly composed of three parts. Firstly, at the top of the page is a search box (**Figure 3**A). Click button named all to switch to other proprietary databases, such as literature, drugs and GSA. Under the search bar is a switch to advanced search, and by clicking advanced search, users enter advanced search. Advanced search is a keyword search on literature, drugs, GSA and patent database. At the bottom of the search page is the statistics of OOCDB. At present, OOCDB collected 122,000 academic papers related to organs-on-a-chip, including more than 4000 academic journals and 20,000 affiliations which involve 25,000 researchers and experts in the field. Secondly, the core parts of our database are popular resources, featured tools and publication statistics (Figure 3B). Popular resources include eight sub databases: GSA, drug bank, device, organs, patents, cancer research, chemicals and toxicants. View datasets takes users to desired sub database and query the corresponding information. Featured tools include three-dimensional model, mathematical modeling and citation map. The user can process the data by entering the corresponding section. Publication statistics includes top authors, top affiliation and top journal. Users can query their details by clicking more. Lastly, As shown in Figure 3C, the page shows the basic information of our database. Such as the resource, feature of database and information about developers.

**Figure 3.**
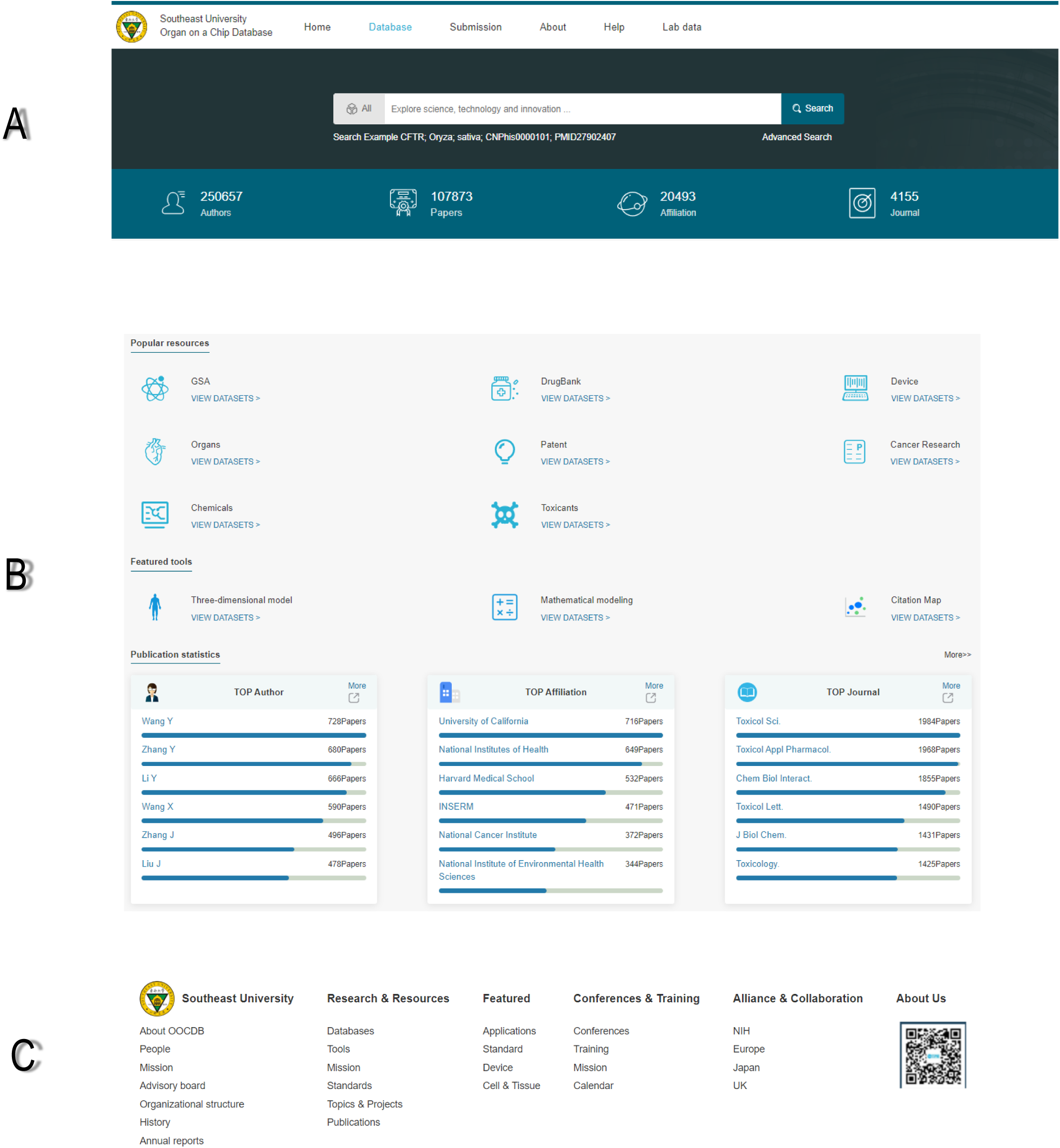
The screenshot of the main interface of OOCDB shows four parts of the website. **A**. The search box of OOCDB and the data introduced. **B**. The main body of the database, including featured tools, popular resources and publication statistics. **C**. The basic information about the OOCDB.

### Popular resources

#### GSA database

A total of 16,846 cases of microarray and transcriptome sequencing raw data were collected in GSA database. These data are categorized based on human organs and tissues types, including lung, muscle, skeleton, nervous system, genetic system, soft tissue and so on. The exact number of microarray and transcriptomic sequencing can be found on database website. In addition, publication years of organs-on-a-chip related literature are tracked. And the statistical results show that in recent years, the research data of organs-on-a-chip is on the rise, and organs-on-a-chip is gradually becoming a trending research area. In OOCDB, data in GSA can be searched using ordinary search and advanced search. Information can be retrieved by keywords when using ordinary search. Advanced search includes and, or, not relationships, which can be searched by multiple keywords with relationships between them. In addition, it also includes access number, title, experience type, organization name, department and lab, which can be searched, respectively. Search result presents information such as status, title, organization(s), experience type, summary, overall design, contributor(s), platforms, samples, and relations. Also, users can find the relevant literature data by clicking the link of the title in the GSA database. These data are stored in the OOCDB and are available for download in soft, miniml, and txt formats. *DrugBank database*

The database of DrugBank contains information of 14,143 kinds of drug related to organs-on-a-chip. The database, as shown in the **Figure 4A**, provides the drug information like name, accession number, structure, chemical formula, synonyms, UNII (Unique Ingredient Identifier), InChi (International Chemical Identifier), InChi key, SMILES (Simplified molecular input line entry system), weight, drug entry, and other drug information, which is convenient for users to find drug-related information. The searching methods of drugs also include ordinary search and advanced search. The advanced search also includes and, or, not relationships, and information can be found by entering drug name, type, CAS (Chemical Abstracts Service) Number, description, UNII or groups. In addition, OOCDB provides literature data on the application of this drug in organs-on-a-chip if this drug is used in organs-on-a-chip-related experiments.

**Figure 4.**
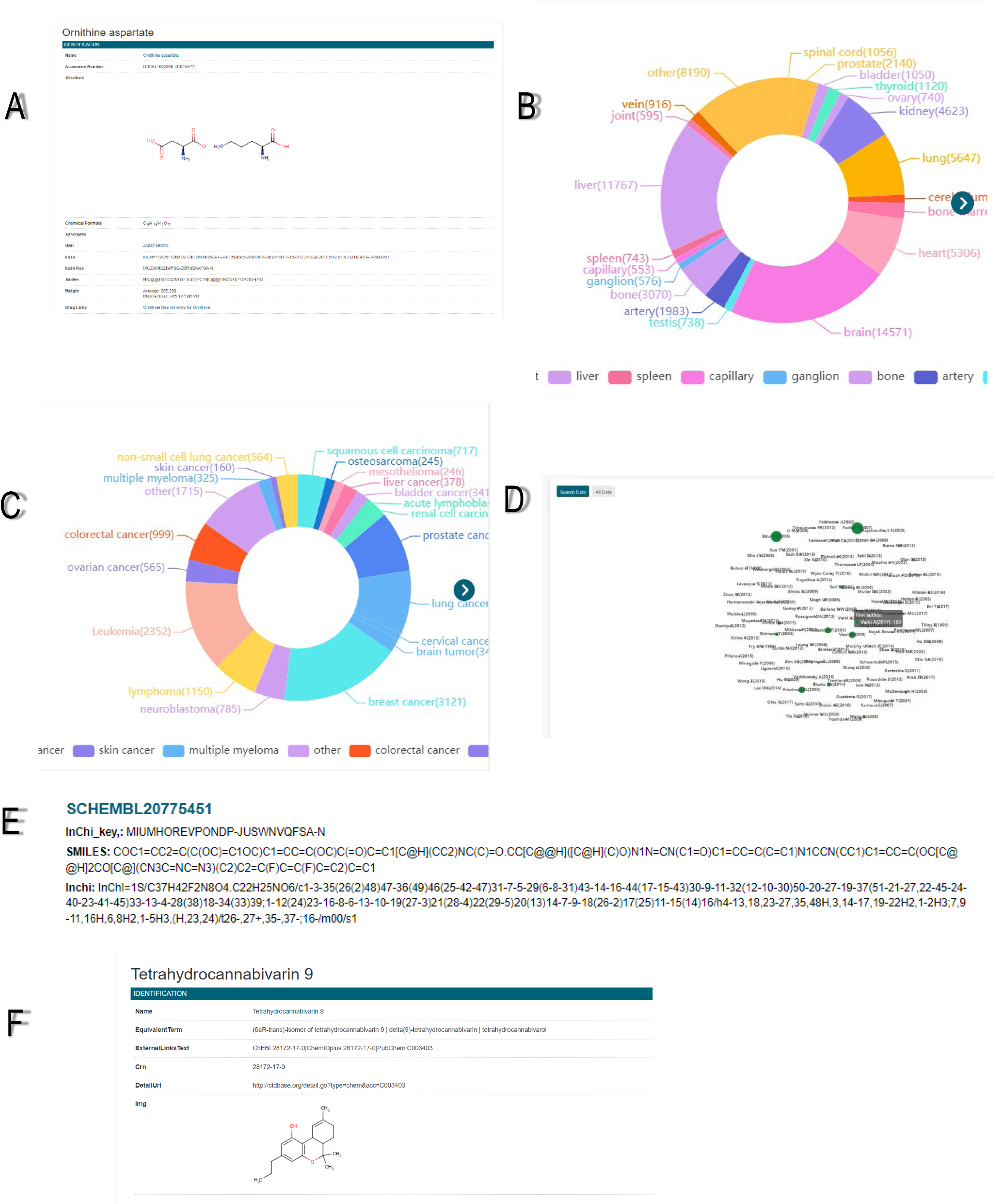
Brief introduction to the sub-databases. **A**. Drug detail interface in the Drugbank database. **B**. Organ distribution map in the Organs Database. **C**. Cancer distribution map in the Cancer Research database. **D**. Citation map of the all data. **E**. Chemical detail interface in the Chemicals database. **F**. Toxicant detail interface in the Toxicants database.

#### Device database

In the device database, we first present the distribution map of 160 literatures about organs-on-a-chip devices (**Table 1**). Users can click desired organs-on-a-chip devices in the distribution map to retrieve the relevant literatures in the literature database. The database contains a brief introduction of organs-on-a-chip devices, including fabrication materials, production technology, fluid drive and control in microfluidic chip and signal detection.

**Table 1.** The number of literatures for each Device in the Device database.

| Device name | Number of literatures |
| --- | --- |
| TCPS | 1 |
| PDMS | 52 |
| Hydrogel | 22 |
| Membrane | 60 |
| 3d print | 1 |
| Microfabrication | 18 |
| Machining | 4 |
| Pouring | 2 |
| **Total** | **160** |

#### Organs database

In the organs database, literatures are classified based on organ names, which resulted in 97 different categories and each category corresponds to an organ. Comparing these category keywords with the organs-on-a-chip literature we downloaded, a total of 67,562 relevant documents were screened (**Table 2**), and then classified these literatures, select the top 20 organs with the highest frequency for graphic display. As shown in the Figure 4B, in the organ database, brain (14,571), liver (11,767) and lung (5647) have the most literatures on organ microarray, which is also the research hotspot of organ microarray. OOCDB provides full-text downloads of these related literatures, and provides maps of references and citations. The reference and citation are drawn into a citation map, and size of the circle indicates the author’s influence in this field. Larger circle represents greater influence of authors, which is convenient for researchers to find experts in the field.

**Table 2.** The number of literatures for each organ in the Organs database.

| Organ name | Number of literatures |
| --- | --- |
| Brain | 14571 |
| Liver | 11767 |
| Lung | 5647 |
| Heart | 5306 |
| Kidney | 4623 |
| Bone | 3070 |
| Prostate | 2140 |
| Artery | 1983 |
| Bone marrow | 1443 |
| Other | 15892 |
| **Total** | **67562** |

#### Patents database

Patent database is developed by screening out 54 key words related to organs-on-a-chip, searching on google patents, and retrieving a total of 18,710 patents. These patents are downloaded to our database and classified according to different countries. As shown in **Table 3**, the United States (5339), China (4101) and Japan (1944) hold a large number of patents related to organs-on-a-chip and have a great advantage in the field of organs-on-a-chip. In addition, OOCDB also provides full text retrieval and download of relevant patents. Users can narrow the search scope according to date and country in the OOCDB, and also retrieve in the advanced search according to title, author, abstracts, description, patent number and country expire date of the patent. So, the users can easily download the patents they need.

**Table 3.** The number of patents for each countries in the patent database.

| Country name | Number of patents |
| --- | --- |
| United States | 6339 |
| China | 6101 |
| Japan | 1944 |
| European Patant Office | 969 |
| WIPO(PCT) | 783 |
| France | 552 |
| USSR-Soviet Union | 548 |
| South Korea | 495 |
| Germany | 414 |
| Australia | 356 |
| Other | 1409 |
| **Total** | **19910** |

#### Cancer research database

Cancer research database is similar to the organs database. By using 87 cancer keywords to filter, 17,921 articles were found, and the top 20 frequent cancer mentioned in paper were presented in a chart (Figure 4C). In the cancer database, breast cancer (3121), leukemia (2352) and lung cancer (1624) are mentioned in high frequency, and organs-on-a-chip are widely used in these cancers (**Table 4**). Like the organs database, this database also contains citation map to find out the authoritative scholars in the field of organs-on-a-chip and cancer (Figure 4D).

**Table 4.** The number of literatures for each cancer in the Cancer research database.

| Country name | Number of patents |
| --- | --- |
| Breast cancer | 3121 |
| Leukemia | 2352 |
| Lung cancer | 1624 |
| Prostate cancer | 1570 |
| Lymphoma | 1150 |
| Colorectal cancer | 999 |
| Neuroblastoma | 785 |
| Squamous cell carcinoma | 717 |
| Ovarian cancer | 565 |
| Non-small cell lung cancer | 564 |
| Other | 4474 |
| **Total** | **17921** |

*Note:* The number of cancer research-related literatures contained in each cancer in the cancer research databas.

#### Chemical database

The chemicals database stores the information of 4,986,258 compounds related to organ microarrays. The compounds are ordered by name. In the compounds database, as shown in Figure 4E, provided information including InChi key, SMILES, InChi. It is convenient for users to find the structures and properties of related compounds.

#### Toxicants database

Toxicants database contains information about 17,070 kinds of toxicants. They are arranged in alphabetical order, which can be retrieved according to the order of the alphabet. Users can also directly import the name of the toxicants in the search box and jump to the toxicants interface. In the toxicants detailed interface (Figure 4F), a variety of toxicants information, including name, equivalent terms, other database numbers, CAS registry number, structure image and external links are provided. If some properties of toxicants cannot be found in OOCDB, users can also click external links to enter the special toxicants database for query. At the same time, we also combine the toxicants database with the organs-on-a-chip literature database. If the toxicant is applied to organs-on-a-chip related research, the corresponding literature is provided below.

### Featured tools

OOCDB’s featured tools present the most important features about organs-on-a-chip and the user can get this data. The feature resource includes three-dimensional model, mathematical modeling and citation map.

#### Three-dimensional model

Three-dimensional model is a tool that we provide to realize the combination of organs-on-a-chip data in OOCDB and human organs at the level of human anatomy. Users can navigate and find the target organ by navigating in 3D anatomical atlas for the human brain, male and female body. A visual windows show the real human anatomical structures (**Figure 5**). This part includes three parts: human brain model, male anatomical model and female anatomical model. As shown in the Figure 5, the database provides high-quality human brain model, male anatomical model and female anatomical model including internal structure using a surgical view. Users can click the image to enter the corresponding model, and then click the image to enter the corresponding organs. In each model, many anatomical layers in the human body or brain are provided for users to navigate. They can enter any organ following anatomical map guidance. In each model, number of types of brain organs, male organs and female organs, are 111, 2984 and 3000, respectively. By clicking the corresponding position of the organ in the model, users can get corresponding organ name, and then by clicking the name of the organ, users can view omics data, literature data and drug data on the organ in OOCDB, so as to find the detailed information about the organs-on-a-chip and the current research status. It is beneficial for researchers to find relevant information, and to promote the development of organs-on-a-chip technology.

**Figure 5.**
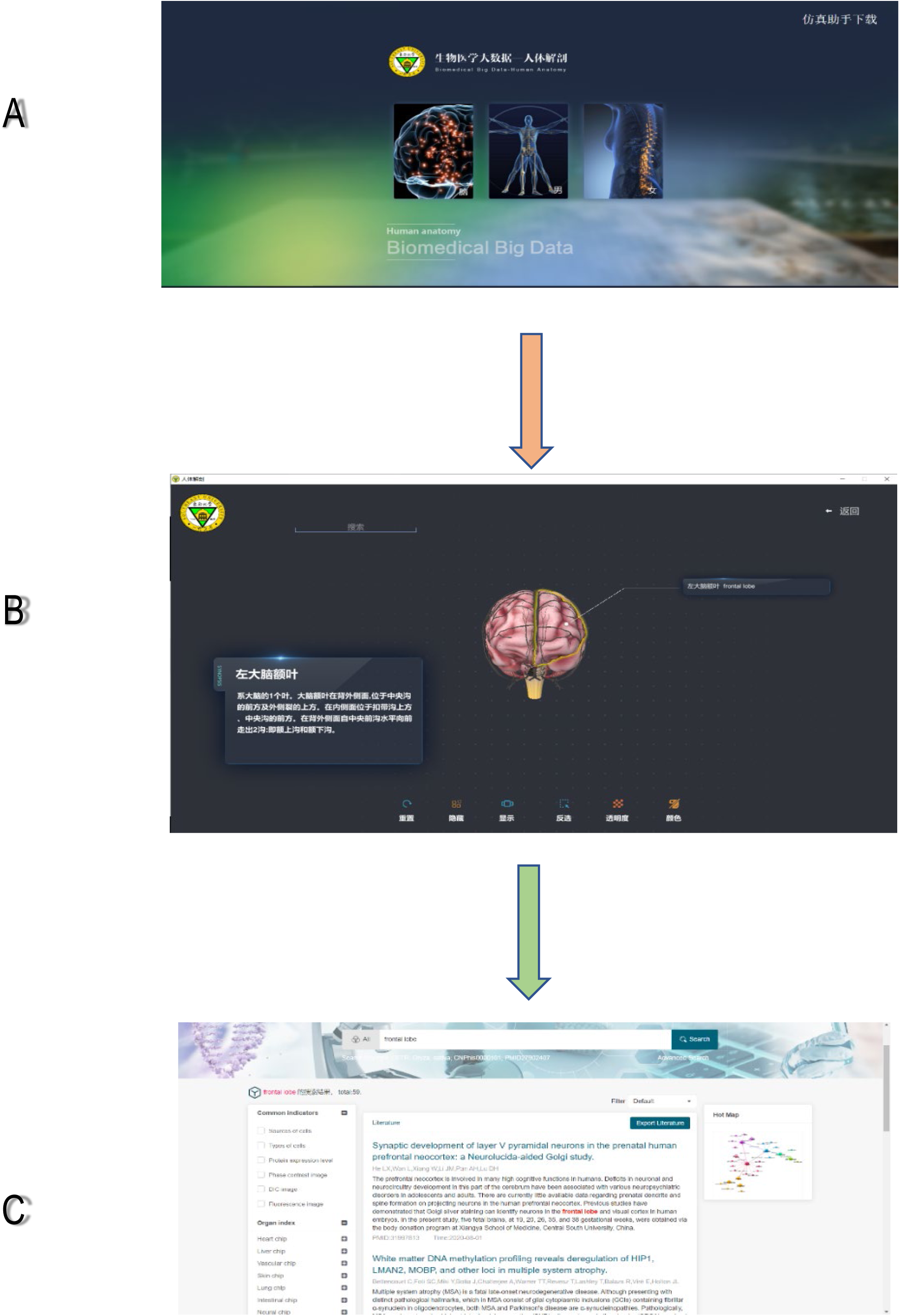
Steps of searching for related literature by the organization on a chip study. **A**. Enter the organization on a chip study sub database. **B**. Enter the 3D model you want to find. **C**. Click on the name of the organ to find the relevant literature.

#### Mathematical modeling

For the most majority of users, their most concerned is whether their own data derived from *in vitro* organs-on-a-chip platform is consistent with the organ in vivo and how much similarity between them. Thus we provide a Mathematical modeling section, which provide users with a useful tool to estimate consistency between *in vitro* data based on their organs-on-a-chip platform and the gene expression data from real human organs. This section currently provides a tissue specific enrichment approach named TissueEnrich, which is a tool to calculate the enrichment of tissue-specific genes in a set of input genes with the hypergeometric test [24]. This tool is embedded to web server with a friendly graphical user interface. User can test the tissue specificity of the cells *in vitro* cultured using organs-on-a-chip platform. The workflow of this tool has three steps: Input, Modeling and Output (**Figure 6A**). For the Input step, user uploads the gene list of the most highly expressed genes, differentially expressed genes, or co-expressed genes to the web server. The gene list can be typed directly or selected as a file in the web page of the tool (Figure 6B). Meanwhile the user can also provide an expression dataset of interest for further tissue specific enrichment analysis in the next step (Figure 6C). After the submit button is clicked by the user, the IDs of the genes uploaded by the user will be checked automatically by a program to confirm that the genes are recognizable for the enrichment analysis. And the users can choose the genes according to the results of the ID conversion as well (Figure 6C). The confirmed genes will be passed to the enrichment analyzer, which will compare the gene list to the prebuilt datasets or the user provided custom expression dataset. To compare to the prebuilt datasets, the user needs to choose whether to use the Benjamini-Hochberg correction for the multiple hypothesis testing, which is recommended to be used. Two prebuilt human datasets are provided for defining the tissue-specific gene expression, i.e. HPA (Human Protein Atlas) [25] and GTEx [26]. If the custom expression dataset is used, the server will direct to the parameters setting page (Figure 6D). After all these setting finished, the server will start to carry out the analysis. The results will be output as tables and plots. The tables are the scores of the enrichment analysis, like the log10(P-value), tissue specific genes number, fold change, samples and tissue type (Figure 6E), and the genes that are enriched (Figure 6F). The plots are the bar plot of the log10(P-value) (Figure 6G), and the heat map of the expression profile of the tissue specific genes (Figure 6H). All the tables and plots are available for downloading for the users.

**Figure 6.**
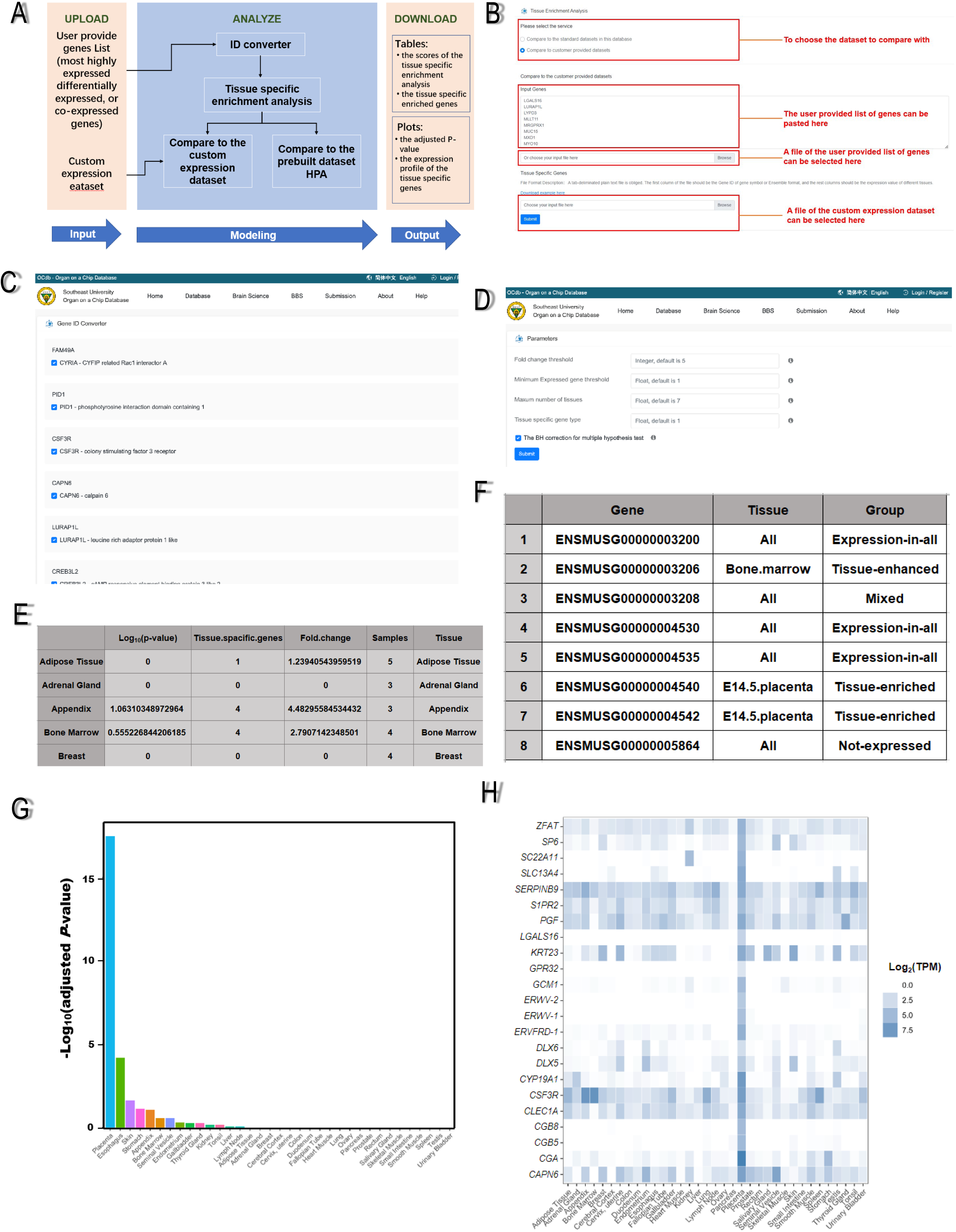
The tissue enrichment analysis was carried out in the mathematical model. **A**. The workflow of the mathematical model includes input, modeling and output. **B**. The input of mathematical model. Firstly, the user chooses the dataset to the compare with. Secondly, the user provided list of genes. Thirdly, the user selected a file of the user provided list of genes. Lastly, the user selected a file of the custom expression dataset. **C**. Tissue specific enrichment analysis. Program checked the genes uploaded by the user to confirm that the genes are recognizable for the enrichment analysis. and the users can choose the genes according to the results of the ID conversion as well. **D**. The parameters setting page. Include Fold change threshold, Minimum Expressed gene threshold, Maxum number of tissues and Tissue specific gene type. **E**. The scores of the enrichment analysis. Include the log10 (P-value), tissue specific genes number, fold change, samples and tissue type. **F**. The genes that are enriched. **G**. The plots are the bar plot of the log10 (P-value). **H**. The heat map of the expression profile of the tissue specific genes.

#### Citation Map

Citation map is a tool used in OOCDB to analyze the citation situation in organ chips. Users can search for a field keyword in citation map’s search box. Citation map will search for authors in the field in the organs-on-a-chip literature database based on the keywords searched by users and present them in a visualized way. In addition, users can click all data to visualize the entire data. As shown in the **Figure 7A**, the size of the circle indicates the number of papers cited by that author in the field, and the larger the circle, the more cited articles the author has in the field. In addition, the citation relationship of each author in the field is also expressed in citation map by connecting the lines, if there is a citation relationship among the authors. Also, a trend graph Figure 7B of publications in the field is given below, showing the change in the number of papers cited in the field of organs-on-a-chip over time. Therefore, the citation map tool can be used to clearly grasp the research hotspots in the direction of organs-on-a-chip and promote the development of organs-on-a-chip related fields.

**Figure 7.**
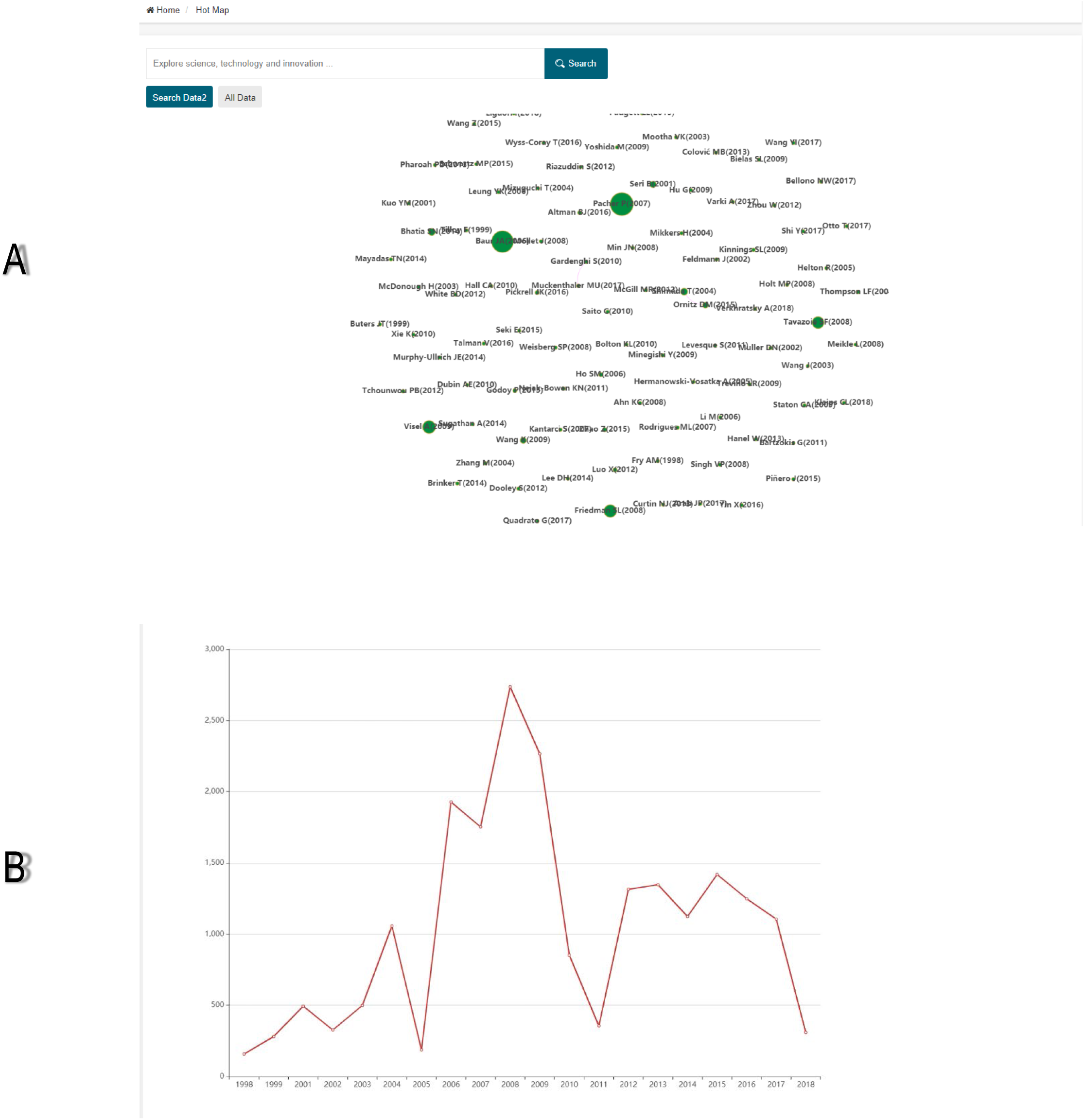
Introduction of citation map. **A**. Graph of authors with a high number of citations in the field. **B**. Change of cited literature in the field over time.

### Lab data

As shown in **Figure 8**, the lab data section includes the data and the model of the organs-on-a-chip from laboratories. At present, it mainly involves the results of our research institute as a demonstration - including blood vessels-on-a-chip, epidermis-on-a-chip, heart-on-a-chip and tumor-on-a-chip and will contains data from our chip’s-end users, collaborators, and other labs’ data (Figure 8A). The model of the organs-on-a-chip section supports search, filtering, and download functions. Users can search directly for relevant organs-on-a-chip data, or filter data by organ and/or by simulation method in order to find the data they expected quickly and easily. At the same time, we supplied the download function which allows users to directly download the data they required (Figure 8B). In addition, users can view detailed organs-on-a-chip data by clicking on “Detail”, which provided detailed information for each organs-on-a-chip. We have divided our own lab data into six fields: name, organ, type, description, device image and model image, so that the data measured/recorded from each chip can be displayed in a well-organized manner for researchers to browse and study in detail (Figure 8C). Furthermore, users can click on device image or model image to view the details of the Image to learn more about the organ chip (Figure 8D).

**Figure 8.**
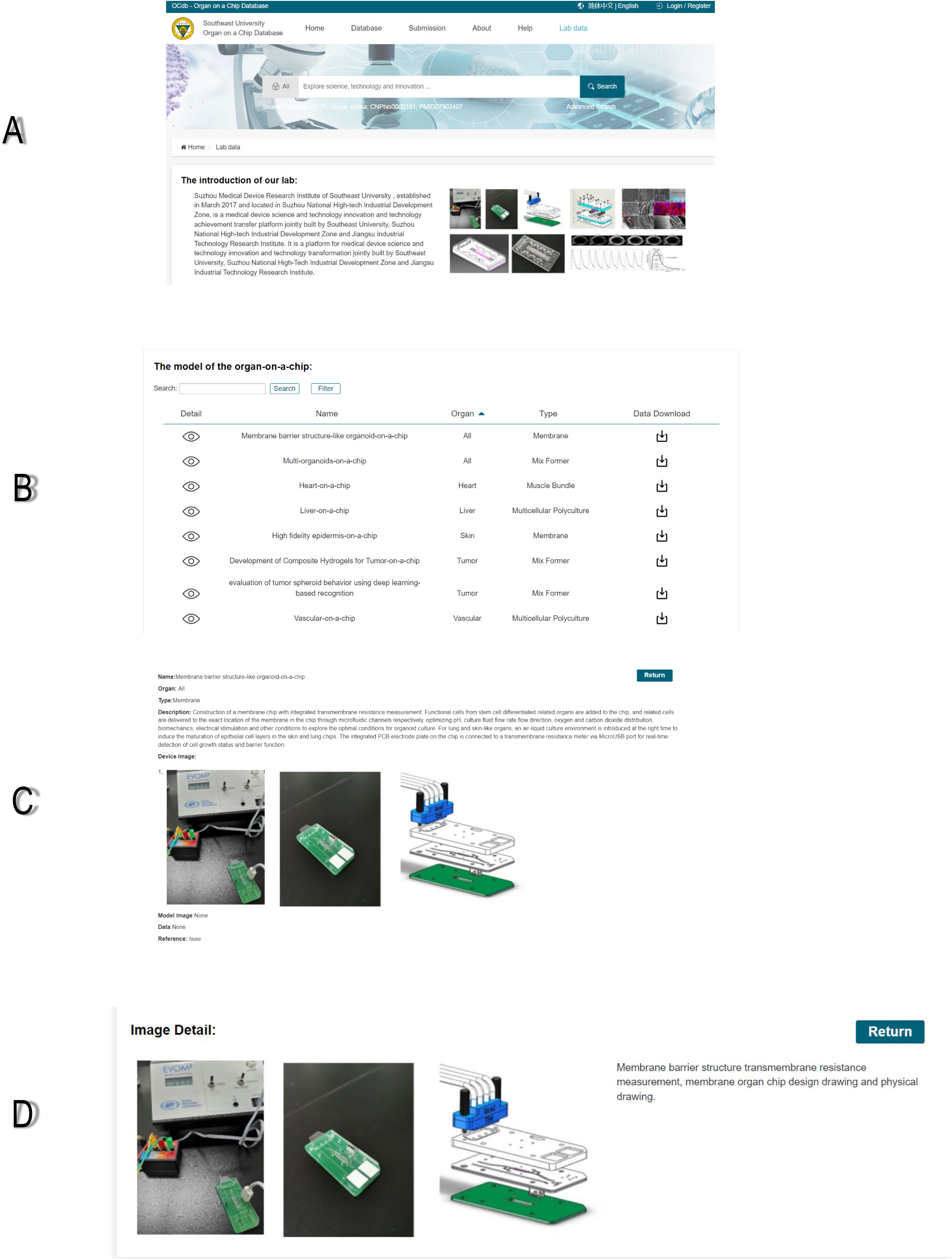
Introduction of our data. **A**. The introduction of organ-on-a-chip data from our lab. **B**. Major organ model data from our lab. **C**. Detailed information on organs-on-a-chip models. **D**. The image details of the organ chip.

### Publication statistics

In addition, OOCDB also collates authors, institutions and journals. Users can select different periods and view the number of published papers, citation frequency and publication trend of top authors, top affiliation and top journals in the current period. Users can sort by count or citation, and choose the authors, institutions, and journals users want to read.

### Case study: using OOCDB to study hepatocellular carcinoma

OOCDB makes it easy and fast to conduct research in the field of organ-on-a-chip. Compared to other open public databases, OOCDB focuses on organ-on-a-chip, making it easy for researchers to find relevant information quickly. Compared with MPS-DB, OOCDB can provide more comprehensive data. The following is an example of what OOCDB can provide for hepatocellular carcinoma.

For the genes associated with hepatocellular carcinoma, we can perform tissue enrichment analysis using a mathematical model with a dataset of tissue-enriched genes from 12 non-tumor tissues [27]. We performed enrichment analysis for these 318 genes using the method in Binghua Li et al [27]. The data was processed by entering the gene symbol and choosing human protein atlas. As shown in Figure 9A, these genes were mainly enriched in the liver, which is the same as the results of Binghua Li et al [27]. These results proved the reliability of the mathematical model. As shown in Figure 9B, the mathematical model highlighted genes that were highly enriched in the liver, such as SERPINA1, APOC1, *etc*. which laid the foundation for subsequent experiments. As shown in Figure 9C, we can also use the 3D model to quickly find where the liver is located in the body and which organs it has adjacent to, and perform a quick search for these organs. OOCDB also provides a citation map function, and in Figure 9D, the results of a citation map search for hepatocellular carcinoma are provided, giving information on experts in the field. For hepatocellular carcinoma, OOCDB provides a large amount of information on literature, histology and drugs (Figure 9E, Figure 9F, Figure 9G), so that researchers can easily and quickly retrieve the information they need to conduct their research.

**Figure 9.**
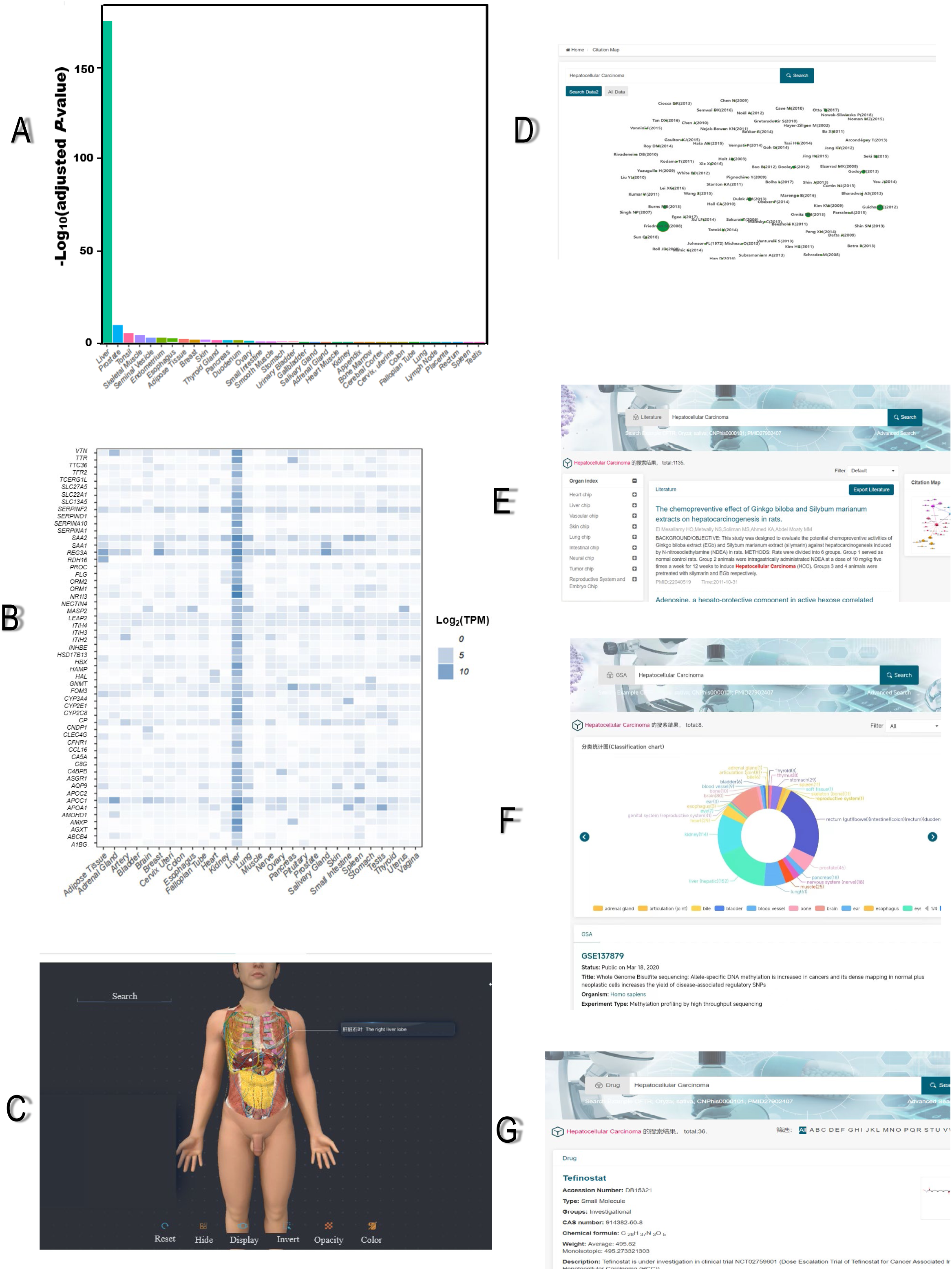
Application of OOCDB in hepatocellular carcinoma research. **A**. Results of tissue enrichment analysis of 318 hepatocellular carcinoma-associated genes. **B**. Heat map of 318 hepatocellular carcinoma-associated genes for specificity analysis. **C**. Three-dimensional model for the localization of the liver. **D**. Search results for hepatocellular carcinoma in citation map. **E**. Search results for hepatocellular carcinoma in the literature database. **F**. Search results for hepatocellular carcinoma in the GSA database. **G**. Search results for hepatocellular carcinoma in the drug database.

### Conclusion and future plans

With the rapid development of modern science and technology, organs-on-a-chip technology has been widely concerned and developed rapidly as soon as it came out. Organs-on-a-chip is a new platform for cell culture *in vitro*, which born from the combination of microfabrication technology and three-dimensional cell culture. Compared with the traditional methods of building models, it has the advantages of portability, high throughput and in vivo microenvironment simulation. It also has broad application prospects in the study of disease pathogenesis and drug screening. It can provide an integrated and systematic solution for life sciences and medical research, and play an important role in many fields such as life science and medicine. Moreover, because organs-on-a-chip technology is a new technology, the data generated by it is currently in the early stage of the “blowout” state. Many related literatures of organs-on-a-chip are constantly published, and the research of organs-on-a-chip is becoming mature. Therefore, it is necessary to establish a database related to organs-on-a-chip.

Organs-on-a-chip database (OOCDB) is a unified platform (science as a service) that provides organs-on-a-chip biological big data sharing and application services for scientific research communities. It is based on big data, artificial intelligence and cloud computing technology provides data services specialized for organs-on-a-chip data archiving, computational analysis, knowledge search, sharing services, and visualization.

In OOCDB, we have done the corresponding data archiving, annotation, analysis and visualization work, and finally formed dozens of sub databases and analysis tools. OOCDB also searches and builds an index, and correlates these data with organ tissue, drug, poison and disease model research, so as to realize the traceability of the entire process of organs-on-a-chip data from specific organization to project research to information data to achieve comprehensive data. We will continue to enrich our synonym database to ensure more synonym searches are supported. In addition, the 3D models and mathematical models in OOCDB are a valuable resource for developers and users who develop and use human organ model predictions, effectively promoting the safety and accuracy of the models. We will continue to translate the content in the 3D model into English. More importantly, OOCDB will be updated every 3-6 months by dedicated staff to complete the latest academic papers and related information. In addition, it will also provide the latest progress in the field of organ-on-a-chip research, analysis of research hotspots, and real-time online statistics in academic related fields. Thus, it will provide the most comprehensive, real-time, and dynamic latest progress of on-chip organ research for global on-chip organ researchers and experts in industry and government health medicine. So our database can promotes the development of organs-on-a-chip field. In addition, due to the serious outbreak of the COVID-19, we will increase the relevant literature collection on lung chip and new crown pneumonia epidemic, and contribute to the common difficulties.

## Data availability

OCDB is available at http://www.organchip.cn.

## CRediT author statement

**Jian Li:** Conceptualization, Methodology, Software, Validation, Investigation, Resources, Writing - Original Draft, Writing - Review & Editing. **Weicheng Liang:** Methodology, Software, Validation, Formal analysis, Investigation, Writing - Original Draft, Writing - Review & Editing. **Zaozao Chen:** Methodology, Validation, Formal analysis, Resources, Data Curation, Writing - Review & Editing. **Xingyv Li:** Investigation, Formal analysis. **Pan Gu:** Validation, Writing - Review & Editing. **Anna Liu:** Validation. **Pin Chen:** Validation. **Qiwei Li:** Resources. **Xueyin Mei:** Investigation. **Jing Yang:** Investigation. **Jun Liu:** Resources. **Lincao Jiang:** Validation. Writing - Review & Editing. **Zhongze Gu:** Conceptualization, Supervision, Writing – Review and Editing, Funding acquisition. All authors read and approved the final manuscript.

## Acknowledgments

This study is mainly supported by the National Natural Science Foundation of China (Grant No. 31871322), the National Key R&D Program of China (Grant No. 2017YFA0700500) and the Fundamental Research Funds for the Central Universities (Grant Nos.2242020k10001 and 2242019k10016).

